# Escaping from multiple visual threats: Modulation of escape responses in Pacific staghorn sculpin (*Leptocottus armatus*)

**DOI:** 10.1101/2021.08.12.456046

**Authors:** Hibiki Kimura, Tilo Pfalzgraff, Marie Levet, Yuuki Kawabata, John F. Steffensen, Jacob L. Johansen, Paolo Domenici

## Abstract

Fish perform rapid escape responses to avoid sudden predatory attacks. During escape responses, fish bend their bodies into a C-shape and quickly turn away from the predator and accelerate. The escape trajectory is determined by the initial turn (Stage 1) and a contralateral bend (Stage 2). Previous studies have used a single threat or model predator as a stimulus. In nature, however, multiple predators may attack from different directions simultaneously or in close succession. It is unknown whether fish are able to change the course of their escape response when startled by multiple stimuli at various time intervals. Pacific staghorn sculpin (*Leptocottus armatus*) were startled with a left and right visual stimulus in close succession. By varying the timing of the second stimulus, we were able to determine when and how a second stimulus could affect the escape response direction. Four treatments were used: a single visual stimulus (control); or two stimuli coming from opposite sides separated by a 0 ms (simultaneous treatment); a 33 ms; or a 83 ms time interval. The 33 ms and 83 ms time intervals were chosen to occur shortly before and after a predicted 60 ms visual escape latency (i.e. during Stage 1). The 0 ms and 33 ms treatments influenced both the escape trajectory and the Stage 1 turning angle, compared to a single stimulation, whereas the 83 ms treatment had no effect on the escape response. We conclude that Pacific staghorn sculpin can modulate their escape response only between stimulation and the onset of the response, but that escape responses are ballistic after the body motion has started.

**SUMMARY STATEMENT:** Using double stimulation from opposite sides at different time intervals to simulate coordinated predatory attacks, Pacific staghorn sculpin escape away from the first stimulus, but were unable to turn away from the second stimulus while the escape response was in progress.

## INTRODUCTION

Fish avoid predators by performing sudden accelerations, i.e. fast start escape responses (Domenici and Blake, 1997). The kinematics and neural control of escape responses have been widely investigated (Domenici and Blake, 1993; Domenici and Hale, 2019; Eaton et al., 2001; Stewart et al., 2014). Fish escape responses typically consist of C- or S-starts, based on the shape of the fish at the end of the first contraction (Domenici and Hale, 2019). In C-starts, fish bend their bodies into a C-shape during the initial muscle contraction (Stage 1) usually away from the threat, while a subsequent return flip of the tail (when present, Domenici and Hale, 2019) can produce further acceleration (Stage 2) (Fleuren et al., 2018). Fish can be startled using a variety of stimuli, from mechano-acoustic to tactile and visual stimuli (Domenici and Hale, 2019). The shortest latencies are typically associated with the stimulation of the mechano-acoustic sensory system leading to the activation of the Mauthner cells (M-cells) (Korn and Faber, 2005), whereas visual stimuli tend to show longer latencies because of the longer neural pathway (Mirjany and Faber, 2011) from the optic nerve to the M-cell via the optic tectum (Temizer et al., 2015). Mauthner cell ablation was shown to delay the escape response and to decrease survival in predator–prey encounters (Hecker et al., 2020). Most previous studies on fish escape responses have focused on a single threat such as a model or a real predator approaching, resulting in an escape response directed away from the threat (Domenici and Blake, 1993; Stewart et al., 2013; Stewart et al., 2014; Walker et al., 2005). However, in nature, multiple predators may attack prey from two or more directions simultaneously or in close succession (Amo et al., 2004; Bshary et al., 2006; Stander, 1992; Steinegger et al., 2018). Multiple co-occurring threats are known to affect the prey’s escape directions; for example, lizards escape at ∼180 degree away from single predators but at perpendicular to the predators when attacked simultaneously from two opposite directions (Cooper et al., 2007).

Previous work has investigated the possibility that a modification of the escape trajectory can occur after initial stimulation. Importantly, inhibition of the mechanosensory input occurs during Stage 1 in both C- and S-starts (Russell, 1976), leading Eaton et al. (1981) and Eaton and Emberley (1991) to suggest that the neural command underlying the escape response is ballistic once the movement has begun (i.e. without further sensory information to compute its trajectory). Indeed, Eaton et al. (1988) found that the Stage 2 command of the goldfish is preprogrammed and not dependent on sensory feedback, however it remains unknown if sensory feedback can occur before or after the initiation of Stage 1.

Some taxa are able to modulate escape responses to multiple successive attacks. Certain crickets, for instance, were found to use two escape modes (i.e. running and jumping) with different degrees of flexibility: when crickets escape through running from an initial predator attack, they were able to modulate their trajectory in response to a second attack. However, this was not that case for crickets which escaping by jumping from the initial attack (i.e. a ballistic response) (Sato et al., 2019). In fish, recent work has shown that larval zebrafish may be able to integrate sensory information from multiple threats during delayed escape responses due to a cluster of 38 prepontine neurons that are not part of the fast escape neural pathway (Marquart et al., 2019). This finding suggests that during the initial escape latency (i.e. before the onset of Stage 1 contraction), fish may have the potential to integrate sensory information from multiple threats. Additionally, Domenici and Blake (1993) suggested that sensory feedback may occur after the onset of Stage 1, resulting in a correction of escape trajectories during Stage 2. Hence, the extent of Stage 2 may, at least in part, be controlled by a feedback system (Domenici and Blake, 1993).

Here, we investigated the possibility that escape kinematics may vary depending on the time difference between the two visual threat stimuli coming from opposite sides. Visual looming stimuli are known to trigger an escape response once a given threshold (that depends on the size and speed of the approaching object) is reached (Cade et al., 2020; Hein et al., 2018). We hypothesize that if the escape response is fully ballistic from the time of the first stimulation, escape kinematics will not be modified by a second stimulus delivered at any time interval > 0 ms after the first one. If, in contrast, escape kinematics is modified by a second stimulus, this indicates that sensory feedback is possible during that time interval.

## MATERIALS AND METHODS

### Ethics statement

All animal care and experimental protocols followed the guidelines of the Institutional Animal Care and Use Committee at the University of Washington, Seattle, WA, USA (Protocol No. 4238-03).

### Model species and housing conditions

Pacific staghorn sculpin [*Leptocottus armatus*; 13.9 ± 1.71 cm total length (TL); mean ± standard deviation (s.d.); *n* = 71] were captured by beach seining at Jackson Beach, south of San Juan Island, Washington, USA (48°31’11” N, 123°0’45” W) in July 2019. The fish were maintained in two acrylic tanks (87 cm length × 57 cm width × 14 cm depth) with flow-through seawater under a 14/10 h light/dark photoperiod, at 12.5 ± 0.5 °C (mean ± s.d.). They were acclimatized for ≥ 24 h and fed shrimp pieces every second day. At the end of the experiment, they were released at Jackson Beach.

### Experimental setup

Experiments were conducted in an acrylic fish tank (125.5 cm length × 57 cm width × 35 cm depth; Fig. 1A) filled with seawater at 12.5 ± 0.5 °C (mean ± s.d.). White plastic panels were placed on the tank walls and bottom. A white plastic panel with grid lines (48 cm × 34 cm) was placed at the bottom center of the tank. Two 300 W halogen lamps were set above the tank to illuminate it. A high-speed camera (640 × 360 pixels, 240 fps; Stylus TG-870; Olympus Corp., Tokyo, Japan) was positioned 110 cm above the tank and recorded the escape response.

**Fig. 1.**
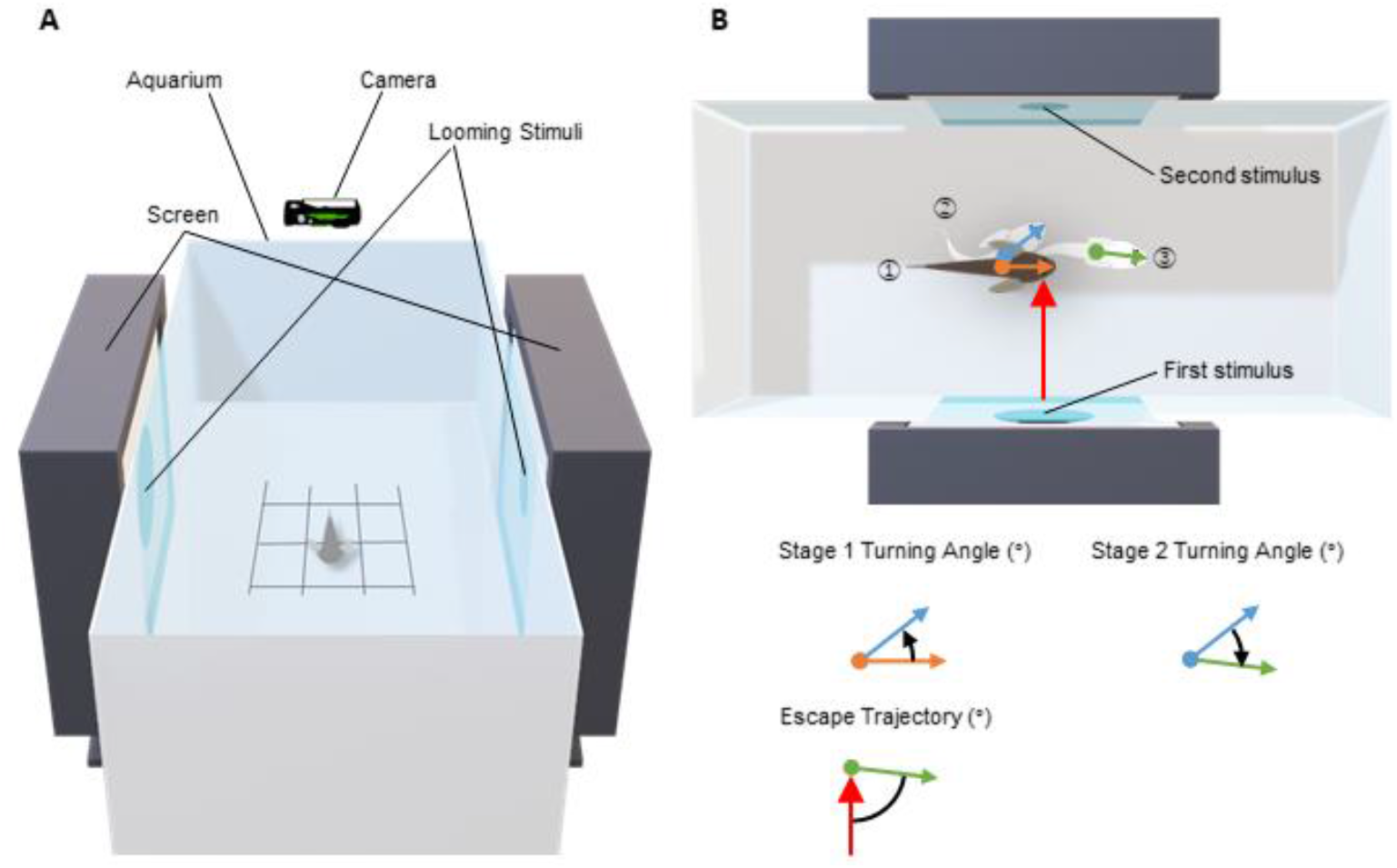
**(A) Schematic representation of experimental setup**. Two screens were used to stimulate fish from the left and right sides with looming stimuli. Fish were always oriented parallel to the long axis of the aquarium (90° initial orientation). **(B) Definitions of Stage 1 and 2 turning angles and escape trajectory**. Upper diagram shows fish just before onset of escape response (1; brown fill), end of Stage 1 (2; white fill) and end of Stage 2 (3; white fill). Lower diagrams show variable definitions. Orange, blue, and green arrows represent fish directions just before onset of the escape response, at end of Stage 1, and end of Stage 2, respectively. Red arrow represents first stimulus direction. Each filled circle of arrows represents fish center of mass (CM).

Two looming stimuli were used. Each stimulus was played on separate screens (1,600 × 1,200 pixels, 60 Hz; DELL 2000FP; Dell Inc., Round Rock, TX, USA) placed centrally on opposing sides of the experimental tank (Fig. 1A). Each stimulus simulated a black disk (24 cm diameter) approaching from 200 cm distances at a constant velocity of 1 m s^-1^. The movie of the looming stimulus (1,600 × 1,200 pixels; 60 fps) was created with R v. 3.6.1 (R Core Team, 2019) using the package *loomeR* v. 0.3.0 (Carey, 2019) and Shotcut Video Editor v. 19.07 (Meltytech LLC, Walnut Creek, CA, USA). For the 0 ms treatment, two identical movies were played simultaneously. For the 33 ms treatment, one of two movies (second stimulus) was played with a delay of two frames (∼33 ms) relative to the first stimulus. For the 83 ms treatment, the second stimulus was played with a delay of five frames (∼83 ms). For the control, only one single looming stimulus was played. The side of the first stimulus was randomized. The times for the delayed stimuli (33 ms and 83 ms) were selected based on the estimated visual escape latency of *L. armatus* (60 ms; Paglianti and Domenici, 2006). Hence, the stimulus delayed by 33 ms was assumed to be within the escape latency of *L. armatus* (defined as the time interval between the stimulus-reaching threshold and the fish response, corresponding to neurosensory processing prior to the visible response; Paglianti and Domenici, 2006) (Fig. 2), whereas the stimulus delayed by 83 ms was assumed to occur during Stage 1 (Fig. 2).

**Fig. 2.**
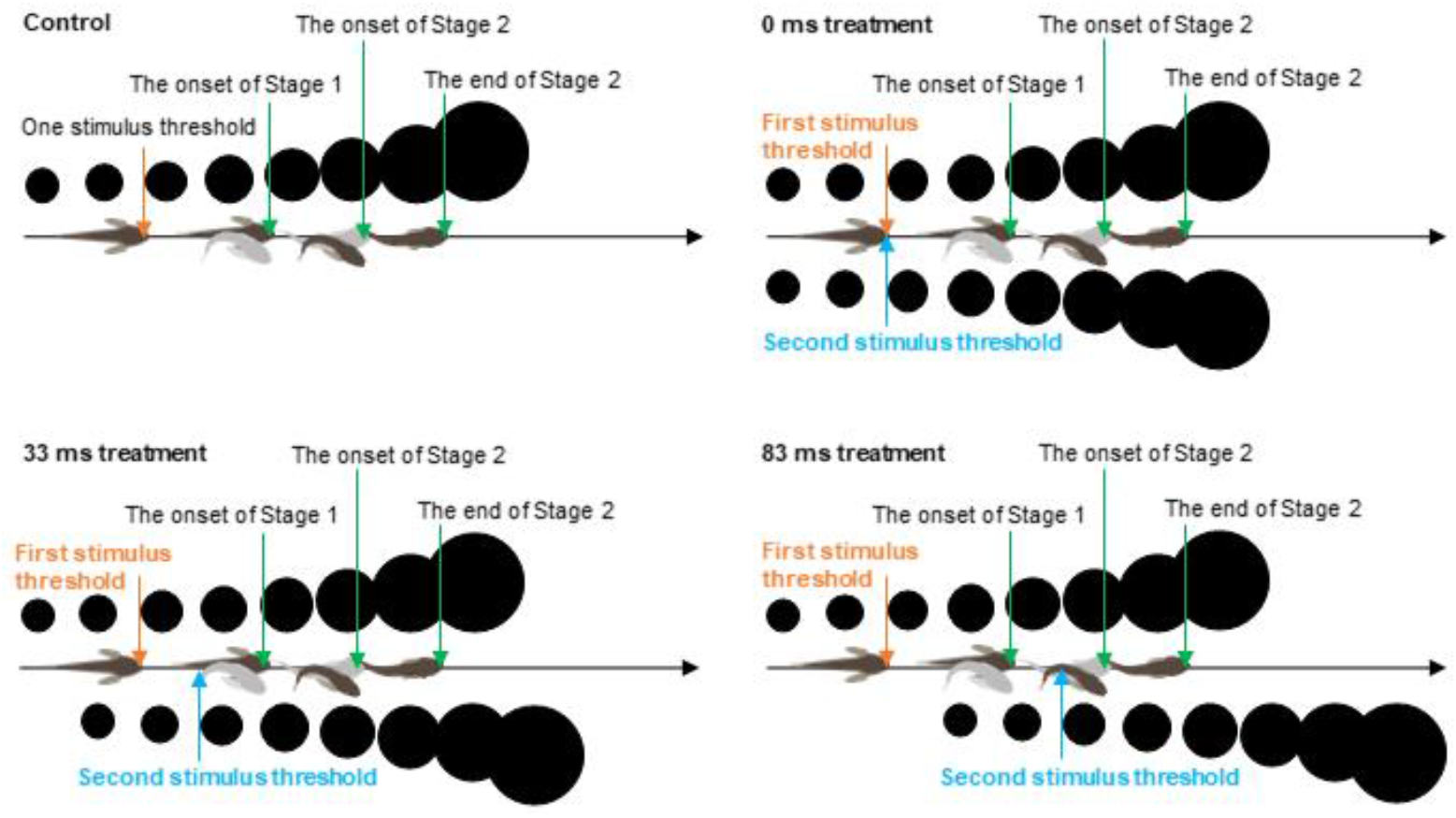
Concepts of the four treatments. Diagrams show transitions of stimuli and fish responses for each treatment. Black disks above are first looming stimuli and disks below are second looming stimuli.

### Experimental procedure

Each fish was transferred to the experimental tank and placed in an opaque PVC shelter (15.5 cm diameter) where it was allowed to acclimatize for 15 min. A square panel under the tank bottom was used as a placement reference ensuring that all fish were placed in the center of the tank at a distance > 1.5 body lengths away from the walls to avoid any interference with their escape trajectory (Eaton and Emberley, 1991). The fish were placed perpendicular to the stimuli by carefully rotating the PVC shelter [86.76° ± 10.93° (mean ± s.d.); *n* = 66].

After acclimatization to the experimental tank, the shelter was removed and fish were left undisturbed for an additional two minutes, after which they were startled. Each fish was exposed to each of the four treatments (in random order) only once. The side of the first looming stimulus was randomly selected. Between stimuli, the fish were returned to the PVC shelter to avoid stimulation prior to each treatment. Fish were allowed over two minutes to recover from the previous stimulation before the next trial continued. If its ventilation was higher than at rest (i.e. a sign of an elevated stress level) by our visual observation, extra time was allocated until the ventilation rate decreased before the next stimulus was played.

### Data and statistical analysis

The 240fps video of the escape response was analyzed frame by frame with Logger Pro v. 3.15 (Vernier Software & Technology, Beaverton, OR, USA). The only responses used were those in which the fish reacted to the stimuli and initiated an escape response to the first stimulus (total 66 responses: 0 ms treatment = 18 responses; 33 ms treatment = 13 responses; 83 ms treatment = 16 responses; control = 19 responses). The fish snout and center of mass [CM; 35% of total body length (Paglianti and Domenici, 2006)] were digitalized in each frame.

A total of six biomechanical and five time-distance variables were then calculated (Dadda et al., 2010; Domenici and Ruxton, 2015). The escape trajectory (°) (variable 1) was calculated as the angle between the direction of the line passing through the CM and the snout at the end of Stage 2 and the virtual movement direction of the first stimulus (Fig. 1A); The Stage 1 turning angle (°) (variable 2) was the angle between the line passing through the CM and the snout at the onset of Stage 1 and the line passing through the CM and the snout at the onset of Stage 2 (Fig. 1B). Stage 1 was taken as the time interval between the onset of Stage 1 and the onset of Stage 2 (Stage 1 turning duration; ms) (variable 3); The Stage 1 turning rate (° s^-1^) (variable 4) was calculated by dividing the Stage 1 turning angle by the Stage 1 turning duration; The Stage 2 turning angle (°) (variable 5) was the angle between the lines passing through the CM and the snout at the onset of Stage 2 and those at the end of Stage 2 (Fig. 1B). Stage 2 was taken as the time from the onset to the end of Stage 2 (Stage 2 turning duration; ms) (variable 6); The Stage 2 turning rate (° s^-1^) (variable 7) was calculated by dividing the Stage 2 turning angle by the Stage 2 turning duration; The apparent looming threshold (ALT; rad s^-1^) (variable 8) triggering the escape response was calculated using the following equation (Eqn. 1) (Dill, 1974):

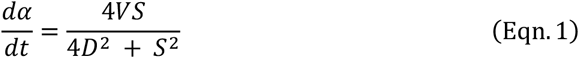

where *D* is the virtual distance between the nearest fish eye and the virtual object (cm), *S* is the size of the virtual object (24 cm virtual diameter), and *V* is the apparent speed of the approaching object (100 cm s^-1^).

The time-distance variables [ maximum acceleration (m s^-2^) (variable 9); maximum speed (cm s^-1^) (variable 10); and cumulative distance (cm) (variable 11)] were measured based on the CM displacement. These variables were evaluated between the onset of Stage 1 and the end of Stage 2. Maximum acceleration and maximum speed were calculated by first- and second-order differentiation, respectively, of the cumulative distance for the time series. A Lanczos five-point quadratic moving regression method (Lanczos, 1956) was applied to calculate these last two values.

To test the difference between treatments, we analyzed each of the 11 variables with an information-theoretic (I-T) approach using Akaike information criterion (AIC), which can be used for multiple comparisons between treatments, and have several advantages over conventional methods such as Tukey HSD (Burnham et al., 2011; Dayton, 1998; Sugiura, 1978). The I-T approach allows comparisons of models with differing distributions (e.g. Gaussian mixture distributions) (Domenici et al., 2008) and nesting/non-nesting (Halsey, 2019; Richards et al., 2011), and are robust to the fact that the distributions of some variables in our data were not unimodal (based on a visual assessment, Fig. S1). Our four treatments (0ms, 33ms, 83ms and control) allowed for 15 combinations of comparisons by categorizing each group as the same (=) or different (≠) (e.g. 0 ms = 83 ms ≠ 33 ms ≠ control, see Table 2 for combination details). These 15 combinations were then analyzed for the best fit using an expectation-maximization (EM) algorithm to fit one-to nine-component Gaussian mixture distributions (GMD) with equal and unequal variance. The EM algorithm constructed 17 GMDs to each of the 15 combinations, equating to 255 models in total, all of which were compared for best fit (see Fig. S2 for the graphical explanation of the I-T approach and estimations of the GMDs). AIC was calculated with the following equation:

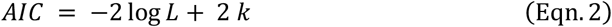

where *k* is the number of parameters and log *L* is the model log-likelihood. *k* was calculated with the following equation:

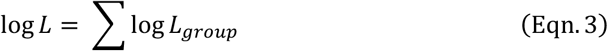

where *k*_*gaussian*_ is the number of parameters of the GMD (see Table 1), *N*_*group*_ is the number of groups in the model (see Table 2).

**Table 1.**
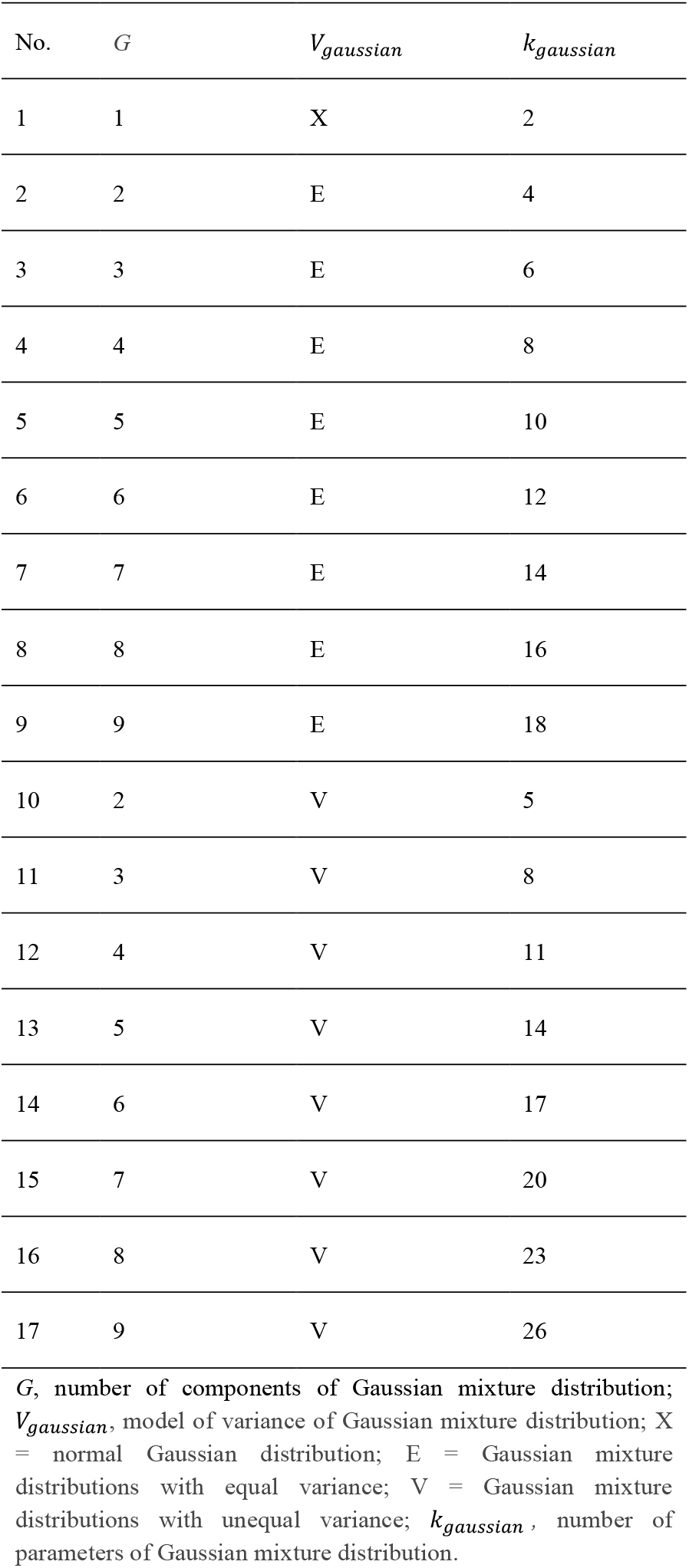
Models of Gaussian mixture distributions.

| No. | $G$ | $V_{\text{gaussian}}$ | $k_{\text{gaussian}}$ |
| --- | --- | --- | --- |
| 1 | 1 | X | 2 |
| 2 | 2 | E | 4 |
| 3 | 3 | E | 6 |
| 4 | 4 | E | 8 |
| 5 | 5 | E | 10 |
| 6 | 6 | E | 12 |
| 7 | 7 | E | 14 |
| 8 | 8 | E | 16 |
| 9 | 9 | E | 18 |
| 10 | 2 | V | 5 |
| 11 | 3 | V | 8 |
| 12 | 4 | V | 11 |
| 13 | 5 | V | 14 |
| 14 | 6 | V | 17 |
| 15 | 7 | V | 20 |
| 16 | 8 | V | 23 |
| 17 | 9 | V | 26 |

$G$ , number of components of Gaussian mixture distribution; $V_{\text{gaussian}}$ , model of variance of Gaussian mixture distribution; X = normal Gaussian distribution; E = Gaussian mixture distributions with equal variance; V = Gaussian mixture distributions with unequal variance; $k_{\text{gaussian}}$ , number of parameters of Gaussian mixture distribution.

**Table 2.** Models used in the information-theoretical approach analysis.

| No. | Model | $N_{group}$ |
| --- | --- | --- |
| 1 | 0 ms = 33 ms = 83 ms = control | 1 |
| 2 | 0 ms = 33 ms = 83 ms ≠ control | 2 |
| 3 | 0 ms = 33 ms = control ≠ 83 ms | 2 |
| 4 | 0 ms = 83 ms = control ≠ 33 ms | 2 |
| 5 | 33 ms = 83 ms = control ≠ 0 ms | 2 |
| 6 | 0 ms = 33 ms ≠ 83 ms = control | 2 |
| 7 | 0 ms = 83 ms ≠ 33 ms = control | 2 |
| 8 | 0 ms = control ≠ 33 ms = 83 ms | 2 |
| 9 | 0 ms = 33 ms ≠ 83 ms ≠ control | 3 |
| 10 | 0 ms = 83 ms ≠ 33 ms ≠ control | 3 |
| 11 | 0 ms = control ≠ 33 ms ≠ 83 ms | 3 |
| 12 | 33 ms = 83 ms ≠ 0 ms ≠ control | 3 |
| 13 | 33 ms = control ≠ 0 ms ≠ 83 ms | 3 |
| 14 | 83 ms = control ≠ 0 ms ≠ 33 ms | 3 |
| 15 | 0 ms ≠ 33 ms ≠ 83 ms ≠ control | 4 |

$N_{group}$ , number of the groups; “0 ms”, “33 ms”, “83 ms” and “control” indicate 0 ms treatment, 33 ms treatment, 83 ms treatment, and control, respectively. “=” assumed that there is no different between left and right treatments and left and right treatments configure one group. “ ≠ ” assumed there is a difference between left and right treatments. For example, “0 ms = 33 ms ≠ 83 ms = control” means that 0 ms treatment is not different from 33 ms treatment whereas the 0 ms treatment are different from 83 ms treatments and control. 33 treatment is different from 83 ms treatment and control. 83 ms treatment is not different from control.

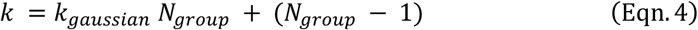

where log *L*_*group*_ is the log-likelihood of each pooled group. The data categorized in the same group were pooled to estimate log *L*_*group*_. For example, in a combination where “0 ms = 33 ms ≠ 83 ms = control”, 0 ms treatment is not different from 33 ms treatment, but is different from 83 ms treatment and control; 33 ms treatment is different from 83 ms treatment and control; 83 ms treatment is not different from control. In that scenario, the data of 0 ms and 33 ms treatments were pooled to estimate the first log *L*_*group*_ and the data of 83 ms and control treatments were pooled to estimate the second log *L*_*group*_. Then, the two pooled log *L*_*group*_ were summed up to calculate the AIC. In an extreme case where “0 ms = 33 ms = 83 ms = control”, data from all four treatments were pooled to estimate the log *L*_*group*_ and AIC, whereas if “0 ms ≠ 33 ms ≠ 83 ms ≠ control”, the data of these four treatments were separately analyzed to estimate each log *L*_*group*_, and the log *L*_*group*_ of four treatments were summed up to calculate the AIC. The most parsimonious model on each of the 11 variables was then chosen based on the lowest AIC. The AIC difference (⊿AIC) was calculated between the best model and all others. Potential models were those with ⊿AIC < 2 (Sugiura, 1978).

All estimations of the GMDs and the analysis of the I-T approach to find differences among treatments were performed in R v. 3.6.1 (R Core Team, 2019) with the *Mclust* v. 5.4.5 package (Scrucca et al., 2016). Because some complex models of each variable could not be calculated with the *Mclust* package due to a singularity in the covariance matrix (Scrucca et al., 2016), the analysis of the I-T approach was performed only on models that could be calculated. Although escape trajectories are circular variables which potentially span 360° (Domenici et al., 2011), most escape trajectories were distributed through a limited arc and the uniformity of the escape trajectories was not supported by Watson’s goodness of fit test for a circular uniform distribution (*U*^*2*^ test; 0 ms: *U*^*2*^ = 0.86, *P* < 0.01, 33 ms: *U*^*2*^ = 0.54, *P* < 0.01, 83 ms: *U*^*2*^ = 0.60, *P* < 0.01, control: *U*^*2*^ = 0.79, *P* < 0.01) and escape trajectories were not distributed around 360° (Fig. 3A). Therefore, the distributions and the difference between treatments were analyzed using linear statistics (estimation of the GMD and the analysis of the I-T approach) as suggested by Batschelet (1981). Calculations of variables and statistical analyses were performed in R v. 3.6.1 with the *circular* v. 0.4-93 package (Agostinelli and Lund, 2017).

**Fig. 3.**
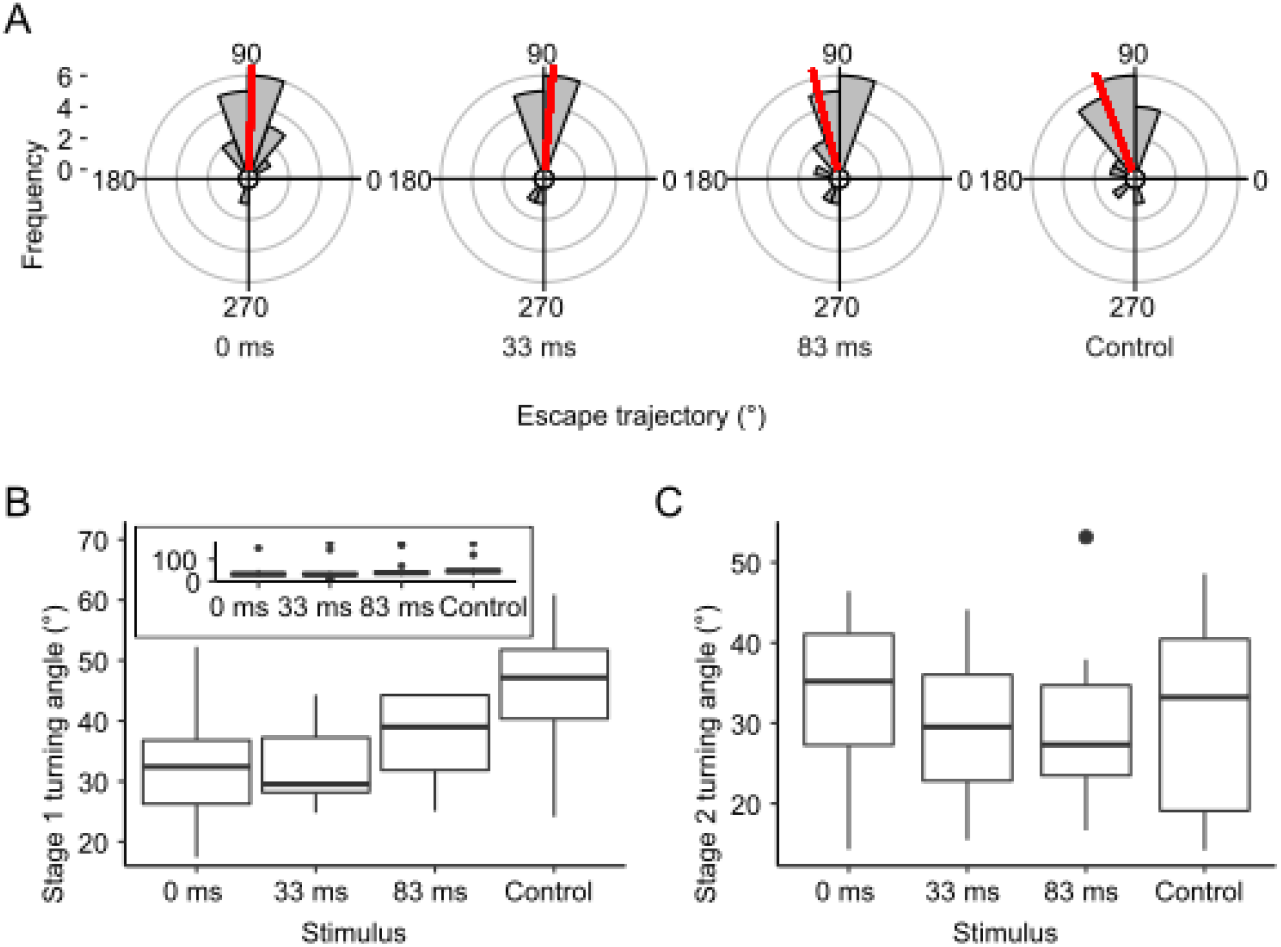
**(A) Circular histogram of escape trajectories**. Red lines show circular mean value of each treatment. The bin intervals are 20°. **(B, C) Boxplots of Stage 1 turning angles excluded outliers (B) and Stage 2 turning angles (C)**. Boxes represent median, lower, and upper quartiles. Ends of vertical lines are minima and maxima. Filled circles are values > 1.5× the upper quartile (outliers). (B) Stage 1 turning angles including outliers is shown in the upper left panel.

## RESULTS

The best models of escape trajectories (variable 1) and Stage 1 turning angle (variable 2, see Table S1) were 0 ms = 33 ms ≠ 83 ms = control” of two-component GMD with equal variance (Table 3 and Figs 3A and 3B). In the escape trajectories, the mean value (*μ*) of the dominant component of the 0 ms and 33 ms treatments’ GMD was lower than that of 83 ms treatment and control’s GMD [0 ms and 33 ms: *μ* = 85.31°; 83 ms and control: *μ* = 104.10°; Table S1]. In the Stage 1 turning angles, the *μ* of the dominant component of 0 ms and 33 ms treatments’ GMD was lower than that of 83 ms treatment and control’s GMD (0 ms and 33 ms: *μ* = 29.90°; 83 ms and control: *μ* = 42.36°; Table S1). The best model for Stage 2 turning angle (variable 5, Table 3) was “0 ms = 33 ms = 83 ms = control”, indicating that it did not differ among the four treatments (Table 3 and Fig. 3C). In the Stage 2 turning rate (variable 7, Table 3), the best model was “0 ms = control ≠ 33 ms = 83 ms” of two-component GMD with unequal variance (Table 3). The *μ* of the dominant component of the 0 ms treatment and the control’s GMD was higher than that of 33 ms and 83 ms treatments’ GMD (0 ms and control: *μ* = 1043.40° s^-1^; 33 ms and 83 ms: *μ* = 586.67° s^-1^; Table S1). In the apparent looming threshold (variable 8), “0 ms = 83 ms= control ≠ 33 ms” of three-component GMD with unequal variance was the best model (Table 3). The *μ* of the dominant component of the 0 ms, 83 ms and control treatments’ GMD was smaller than that of 33 ms treatment’s GMD (0 ms, 83 ms and control: *μ* = 1.60 rad s^-1^; 33 ms: *μ* = 6.88 rad s^-1^; Table S1). Although the best models of the Stage 1 turning rate (variable 4), the Stage 1 turning duration (variable 3) and cumulative distance (variable 11, Table 3) were not “0 ms = 33 ms = 83 ms = control”, ΔAICs of the “0 ms = 33 ms = 83 ms = control” of those variables were < 2 (Table 3). The best models of the remaining three kinematic variables (i.e. Maximum acceleration, Maximum speed and Cumulative distance) were the “0 ms = 33 ms = 83 ms = control”, indicating that there were no differences between treatments (Table 3). The statistical analyses and the summary of the kinematic variables are shown in Tables 2, 3, S1, and S2.

**Table 3.** The results of information-theoretical approach analysis of each variable.

| Variables | Model | $k$ | $\log L$ | $G$ | $V_{\text{gaussian}}$ | AIC | $\Delta\text{AIC}$ |
| --- | --- | --- | --- | --- | --- | --- | --- |
| Escape trajectories (°) | 0 ms = 33 ms ≠ 83 ms = control | 9 | -308.29 | 2 | E | 634.59 | 0.00 |
| Stage 1 turning angle (°) | 0 ms = 33 ms ≠ 83 ms = control | 9 | -272.47 | 2 | E | 562.94 | 0.00 |
| Stage 1 turning duration (ms) | 0 ms = 33 ms = 83 ms ≠ control | 11 | -266.41 | 2 | V | 554.81 | 0.00 |
| | 0 ms = 33 ms ≠ 83 ms = control | 11 | -266.89 | 2 | V | 555.77 | 0.96 |
|  | 0 ms = 33 ms = 83 ms = control | 5 | -272.92 | 2 | V | 555.85 | 1.04 |
| Stage 1 turning rate (° s<sup>-1</sup>) | 0 ms = 33 ms = 83 ms ≠ control | 5 | -489.91 | 1 | X | 989.81 | 0.00 |
| | 0 ms = 33 ms ≠ 83 ms = control | 5 | -490.04 | 1 | X | 990.08 | 0.27 |
|  | 0 ms = 33 ms = 83 ms = control | 2 | -493.69 | 1 | X | 991.38 | 1.57 |
| Stage 2 turning angle (°) | 0 ms = 33 ms = 83 ms = control | 4 | -190.17 | 2 | E | 388.34 | 0.00 |
| Stage 2 turning duration (ms) | 0 ms = 33 ms = 83 ms = control | 4 | -184.40 | 2 | E | 376.79 | 0.00 |
| | 0 ms = 33 ms = control ≠ 83 ms | 13 | -175.82 | 3 | E | 377.64 | 0.85 |
| Stage 2 turning rate (° s<sup>-1</sup>) | 0 ms = control ≠ 33 ms = 83 ms | 11 | -362.48 | 2 | V | 746.96 | 0.00 |
| | 0 ms = control ≠ 33 ms = 83 ms | 9 | -364.88 | 2 | E | 747.76 | 0.80 |
| Apparent looming threshold (rad s<sup>-1</sup>) | 0 ms = 83 ms = control ≠ 33 ms | 17 | -120.80 | 3 | V | 275.61 | 0.00 |
| Maximum acceleration (cm s<sup>-2</sup>) | 0 ms = 33 ms = 83 ms = control | 4 | -592.31 | 2 | E | 1192.61 | 0.00 |
|  | 0 ms = 33 ms = 83 ms = control | 5 | -592.28 | 2 | V | 1194.56 | 1.95 |
| Maximum speed (cm s<sup>-1</sup>) | 0 ms = 33 ms = 83 ms = control | 2 | -317.28 | 1 | X | 638.57 | 0.00 |
|  | 0 ms = 33 ms = 83 ms = control | 5 | -314.35 | 2 | V | 638.70 | 0.13 |
|  | 0 ms = 33 ms = 83 ms = control | 4 | -315.43 | 2 | E | 638.85 | 0.29 |
| Cumulative distance (cm) | 0 ms = 33 ms = 83 ms ≠ control | 11 | -108.08 | 2 | V | 238.17 | 0.00 |
|  | 0 ms = 33 ms = 83 ms = control | 2 | -117.72 | 1 | X | 239.44 | 1.27 |

$k$ , number of parameters of the model; $\log L$ , log likelihood of the model; $G$ , number of components of Gaussian mixture distribution; $V_{\text{gaussian}}$ , model of variance of Gaussian mixture distribution; X = normal Gaussian distribution; E = Gaussian mixture distributions with equal variance; V = Gaussian mixture distributions with unequal variance; AIC, Akaike's information criterion; $\Delta\text{AIC}$ , the difference in AIC between each model and the best model. "0 ms", "33 ms", "83 ms" and "control" show 0 ms treatment, 33 ms treatment, 83 ms treatment, and control, respectively. "=" indicates that there is no difference between left and right treatments. " ≠ " indicates there is a difference between left and right treatments. The best model has a $\Delta\text{AIC}$ value with zero. Each row only shows the results for $\Delta\text{AIC} < 2$ .

## DISCUSSION

The escape trajectories of the staghorn sculpin differed between a single and dual threat stimuli when the fish were visually stimulated from the left and right sides within 33 ms of one another. The mean of the escape trajectories for a single threat stimulus in control was 104.10° (i.e. escaping away from a single threat stimulus). On the other hand, the escape trajectories for dual threat stimuli in the 0 and 33 ms treatments were nearly 90° (i.e. perpendicular to the line of attack of dual threat stimuli). Hence, when attacked from two sides simultaneously or with a short delay (33 ms) between dual threat stimuli, fish tended to escape along a “compromise” trajectory at a similar angle from both stimuli. However, the escape trajectories when the fish were attacked from two sides with a long delay (83 ms) between dual threat stimuli (i.e. when the delayed stimulus occurred after Stage 1 initiation) did not differ from the single stimulus treatment. Consequently, our findings suggest that the escape response of staghorn sculpin is not fully ballistic and sensory feedback may occur during the initial latency period of the escape responses. Once Stage 1 of the escape response is initiated, the escape trajectory is set and is not affected by further feedback control.

Behavioral and neurobiological studies have shown that the Stage 1 turning angles are affected by stimulus direction (Domenici and Blake, 1993; Eaton and Emberley, 1991; Kimura and Kawabata, 2018). The escape response typically avoid predation when the prey evades in the direction opposite to that of the approaching predators (Walker et al., 2005). Hence, Stage 1 turning angles tend to be wide when the stimulus approaches fronting the prey and narrow when the stimulus approaches behind the prey (Domenici and Blake, 1993; Eaton and Emberley, 1991; Kimura and Kawabata, 2018). The optimal escape trajectory was suggested to be between 90° and 180° depending on predator speed (Domenici, 2002; Weihs and Webb, 1984). Here, the fish were stimulated from the left and right sides. Thus, the resulting escape strategy consisted of remaining at an equal distance from both threats by escaping at 90° when the two stimuli were simultaneous. Moreover, as the initial prey orientation relative to the threat was 90°, the fish had to minimize their Stage 1 turning angles to evade using 90° escape trajectories. The results of the present study suggest that the fish minimize their Stage 1 turning angles when they are being attacked from the left and right sides simultaneously.

The mechanism allowing the modulation of Stage 1 turning angles in the 0 ms and 33 ms treatments could be related to the activity of Mauthner cells and associated neurons, as well as the prepontine neurons (Marquart et al., 2019). Prepontine neurons facilitate the integration of multiple sensory information (visual and auditory) and alter Stage 1 of the escape response (Marquart et al., 2019). It is possible that a similar mechanism may occur in the presence of two visual stimuli; i.e. inputs from both eyes might be integrated by prepontine neurons before the onset of escape response. Additionally, the apparent looming threshold of 33 ms treatment was higher than the control, which corresponds to approximately 57 ms increase in the escape latency in the 33 ms treatment. This result suggests that when the fish perceives the second stimulus during the neural processing, it may delay the process based on the single stimulus and integrate the second stimulus information, resulting in 90° escape trajectory from both stimuli.

There were no differences among treatments in terms of their Stage 2 turning angles. In Stage 2, contraction of the body trunk muscles flips the tailfin to the opposite side (Foreman and Eaton, 1993). As acceleration increases during Stage 2, the body slightly rotates, and the final escape direction is determined. The accelerations and propulsive forces and jets are stronger in Stage 2 than they are in Stage 1 (Fleuren et al., 2018; Tytell and Lauder, 2008; Voesenek et al., 2019), and they differ in terms of the relative importance of the rotation or acceleration (propulsion) to their movement (Domenici and Blake, 1993; Domenici et al., 2004; Eaton et al., 1977; Eaton and Hackett, 1984; Tytell and Lauder, 2008; Weihs, 1973). Escape trajectories are related to Stage 1 turning angles (Domenici and Hale, 2019) and the Stage 2 turning angles and rates are smaller than those of Stage 1 (Fleuren et al., 2018; Voesenek et al., 2019). Thus, Stage 2 plays a relatively more important role in acceleration than it does in escape trajectory. The limited effect of stage 2 on the escape trajectory is in line with the lack of differences in Stage 2 turning angles among treatments. Differences between treatments and control were found in cumulative distances, Stage 1 turning durations and rates and the Stage 2 turning rates. These may be related to the differences of the Stage 1 and 2 turning angles.

When the second stimulus reached its threshold after the initiation of the escape response (i.e. in the 83 ms treatment), there was no change in the escape trajectory compared to the control. This is in line with a study on fathead minnows attempting to escape tentacled snakes (Catania, 2009). When fathead minnows were at a strike distance, the tentacled snakes generated a water flow with their bodies and induced a C-start in the fish, directed away from their body but into the snake’s mouth. Prey fish cannot modify their escape response after their reaction to the body-generated water flow because one of two Mauthner cells, which fires first, stimulates the body trunk muscle to initiate an escape response but inhibits activation of the opposite body trunk muscle (Faber et al., 1991; Korn and Faber, 2005). In the 83 ms treatment here, it is likely that the Mauthner cell on the stimulus side was activated, with feedback inhibition preventing the activation of the opposing Mauthner cell (Korn and Faber, 2005) as the latter would result in poor escape response performance. As a result, the 83 ms treatment did not differ from the control. However, our 33 ms treatment suggests that, if stimulated during the neural processing of the first stimulus (i.e. during the escape latency), fish can modify their escape trajectory. Hence, our results demonstrate that fish are capable of receiving additional sensory information during the neurosensory process that involves the circuitry from the optic tectum to the Mauthner cells (Zottoli et al., 1987).

In conclusion, we suggest that the escape response consists of a flexible phase (from the stimulation until onset of Stage 1) where sensory feedback is possible, and a ballistic phase (from the onset of Stage 1 onwards). The ballistic phase is likely to occur in fish fast escape responses as a result of inhibition of the Mauthner cell to trigger a further contraction during stage 1 (Faber et al., 1991). Specifically, a feed-forward inhibitory network guarantees that when one Mauthner cell is excited, (1) it only generates a single action potential (preventing repetitive firing of one Mauthner cell) and (2) the contralateral Mauthner cell will not be activated. As suggested by Faber et al. (1991), this prevents the occurrence of ineffective escape behaviors, characterized by multiple fast body bends, and bilateral muscle contraction which would lead to minimal displacement of the fish. While such ineffective motion patterns are prevented, so is the integration of multiple threats within the time interval that corresponds to stage 1. Hence, effective escape is ensured at the cost of eliminating the flexibility that would be associated with sensory feedback during the early phase of the escape response.

We found that Pacific staghorn sculpin can modulate their escape response only between stimulation and the onset of the response, but that escape responses are ballistic after the body motion has started. Although a flexible phase of Pacific staghorn sculpin lasts for at least 33 ms after the onset of the response, other fish species may show a different relative timing of the flexible and ballistic phases. Identifying which patterns are employed and during which phase, may depend on the species and a phylogenetical analysis of escape flexibility would help us understand the evolution of fish escape response patterns. Furthermore, future research integrating behavioral experiments with neurophysiological measures (e.g. calcium imaging) could allow to understand how the behavioral patterns of the escape response are related to the neural activity when fish are startled by multiple threats.

## ACKNOWLEDGEMENTS

We would like to thank the 2019 Fish Swimming course for support and assistance with fish collections, Friday Harbor Laboratories, University of Washington, for facilities and technical support. H.K. would like to thank his sponsor, Japan/U.S. – E.S. Morse Scholar Exchange Program and M.L. would like to thank the ‘Adopt a Student’ program, the Dudley Fund, and the GRIL (Interuniversity Research Group in Limnology and Aquatic Environments) for their financial support to attend the 2019 Fish Swimming course.

## DATA AVAILABILITY

The datasets generated and used in this study are available from the corresponding authors on reasonable request.

## AUTHOR CONTRIBUTIONS

Conceptualization: P.D.; Methodology: P.D., J.F.S., J.LJ.; Software; H.K.; Formal analysis; H.K., Y.K., T.P., M.L., P.D.; Investigation: H.K., T.P., M.L.; Resources: H.K., T.P., M.L., P.D., J.F.S., J.L.J.; Data Curation: H.K., T.P., M.L.; Writing – Original Draft: H.K.; Writing – Review & Editing: H.K., P.D., J.L.J., Y.K., M.L., T.P.; Visualization: H.K.; Supervision: P.D., Y.K.; Project administration: P.D.

## ABBREVIATIONS

AIC: Akaike’s Information Criterion
ALT: apparent looming threshold
ANOVA: analysis of variance
CM: center of mass
EM: expectation-maximization
GMD: Gaussian mixture distribution
I-T: information-theoretical
SD: standard deviation

## TABLES

**Table S1.**
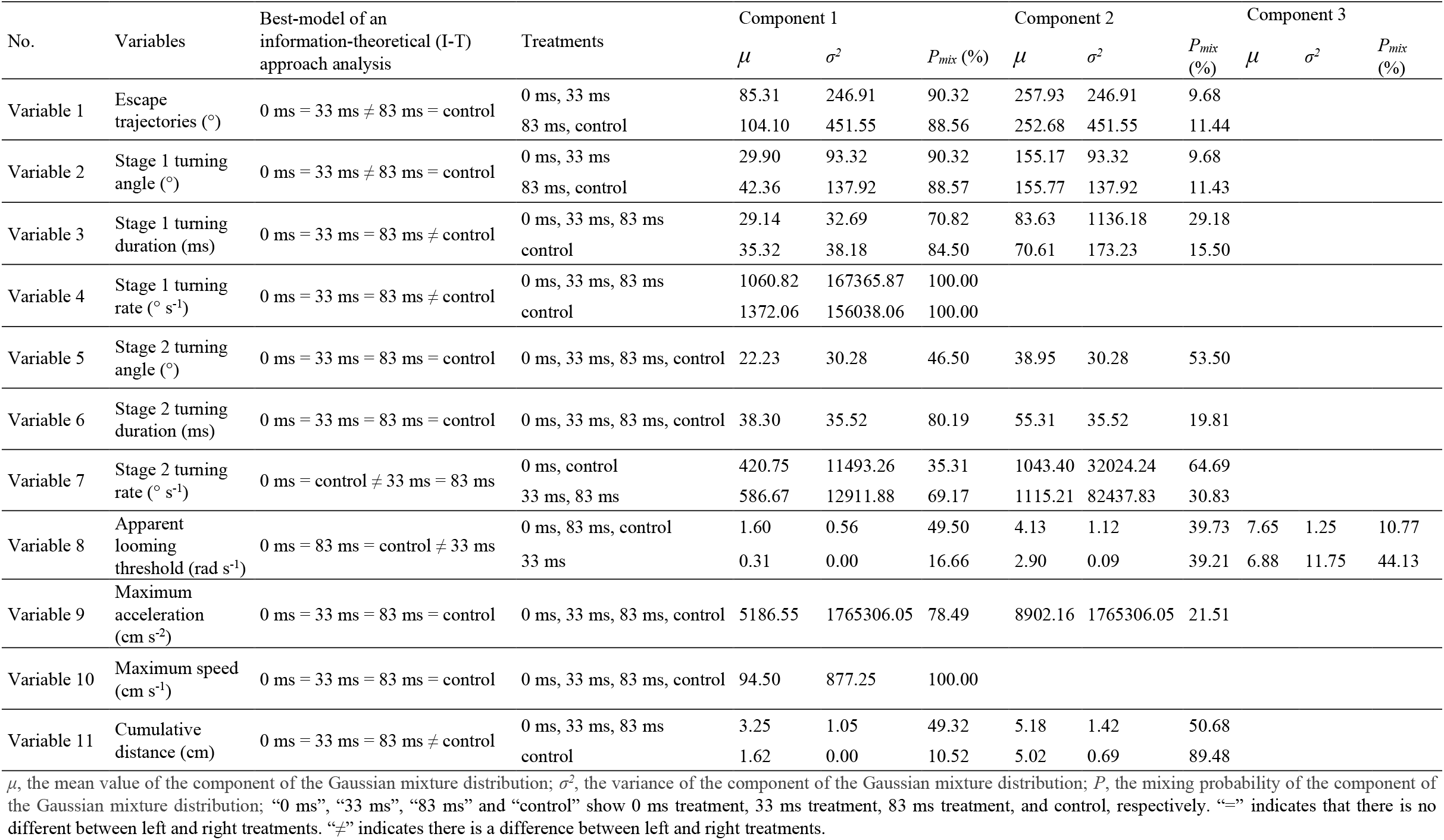
Summary of best fitted Gaussian mixture distributions.

| No. | Variables | Best-model of an information-theoretical (I-T) approach analysis | Treatments | Component 1 |  |  | Component 2 |  |  | Component 3 |  |  |
| --- | --- | --- | --- | --- | --- | --- | --- | --- | --- | --- | --- | --- |
| | | | | $\mu$ | $\sigma^2$ | $P_{mix}$ (%) | $\mu$ | $\sigma^2$ | $P_{mix}$ (%) | $\mu$ | $\sigma^2$ | $P_{mix}$ (%) |
| Variable 1 | Escape trajectories (°) | 0 ms = 33 ms ≠ 83 ms = control | 0 ms, 33 ms | 85.31 | 246.91 | 90.32 | 257.93 | 246.91 | 9.68 | | | |
|  |  |  | 83 ms, control | 104.10 | 451.55 | 88.56 | 252.68 | 451.55 | 11.44 |  |  |  |
| Variable 2 | Stage 1 turning angle (°) | 0 ms = 33 ms ≠ 83 ms = control | 0 ms, 33 ms | 29.90 | 93.32 | 90.32 | 155.17 | 93.32 | 9.68 | | | |
|  |  |  | 83 ms, control | 42.36 | 137.92 | 88.57 | 155.77 | 137.92 | 11.43 |  |  |  |
| Variable 3 | Stage 1 turning duration (ms) | 0 ms = 33 ms = 83 ms ≠ control | 0 ms, 33 ms, 83 ms | 29.14 | 32.69 | 70.82 | 83.63 | 1136.18 | 29.18 | | | |
|  |  |  | control | 35.32 | 38.18 | 84.50 | 70.61 | 173.23 | 15.50 |  |  |  |
| Variable 4 | Stage 1 turning rate (° s<sup>-1</sup>) | 0 ms = 33 ms = 83 ms ≠ control | 0 ms, 33 ms, 83 ms | 1060.82 | 167365.87 | 100.00 | | | | | | |
|  |  |  | control | 1372.06 | 156038.06 | 100.00 |  |  |  |  |  |  |
| Variable 5 | Stage 2 turning angle (°) | 0 ms = 33 ms = 83 ms = control | 0 ms, 33 ms, 83 ms, control | 22.23 | 30.28 | 46.50 | 38.95 | 30.28 | 53.50 |  |  |  |
| Variable 6 | Stage 2 turning duration (ms) | 0 ms = 33 ms = 83 ms = control | 0 ms, 33 ms, 83 ms, control | 38.30 | 35.52 | 80.19 | 55.31 | 35.52 | 19.81 |  |  |  |
| Variable 7 | Stage 2 turning rate (° s<sup>-1</sup>) | 0 ms = control ≠ 33 ms = 83 ms | 0 ms, control | 420.75 | 11493.26 | 35.31 | 1043.40 | 32024.24 | 64.69 | | | |
|  |  |  | 33 ms, 83 ms | 586.67 | 12911.88 | 69.17 | 1115.21 | 82437.83 | 30.83 |  |  |  |
| Variable 8 | Apparent looming threshold (rad s<sup>-1</sup>) | 0 ms = 83 ms = control ≠ 33 ms | 0 ms, 83 ms, control | 1.60 | 0.56 | 49.50 | 4.13 | 1.12 | 39.73 | 7.65 | 1.25 | 10.77 |
|  |  |  | 33 ms | 0.31 | 0.00 | 16.66 | 2.90 | 0.09 | 39.21 | 6.88 | 11.75 | 44.13 |
| Variable 9 | Maximum acceleration (cm s<sup>-2</sup>) | 0 ms = 33 ms = 83 ms = control | 0 ms, 33 ms, 83 ms, control | 5186.55 | 1765306.05 | 78.49 | 8902.16 | 1765306.05 | 21.51 |  |  |  |
| Variable 10 | Maximum speed (cm s<sup>-1</sup>) | 0 ms = 33 ms = 83 ms = control | 0 ms, 33 ms, 83 ms, control | 94.50 | 877.25 | 100.00 |  |  |  |  |  |  |
| Variable 11 | Cumulative distance (cm) | 0 ms = 33 ms = 83 ms ≠ control | 0 ms, 33 ms, 83 ms | 3.25 | 1.05 | 49.32 | 5.18 | 1.42 | 50.68 | | | |
|  |  |  | control | 1.62 | 0.00 | 10.52 | 5.02 | 0.69 | 89.48 |  |  |  |

$\mu$ , the mean value of the component of the Gaussian mixture distribution; $\sigma^2$ , the variance of the component of the Gaussian mixture distribution; $P$ , the mixing probability of the component of the Gaussian mixture distribution; “0 ms”, “33 ms”, “83 ms” and “control” show 0 ms treatment, 33 ms treatment, 83 ms treatment, and control, respectively. “=” indicates that there is no different between left and right treatments. “ ≠ ” indicates there is a difference between left and right treatments.

**Table S2.** Summary of the variables of each treatment.

| Variables | 0 ms | 33 ms | 83 ms | control |
| --- | --- | --- | --- | --- |
| Escape trajectories (°) | 85.29 ± 0.60 (*n* = 18) | 87.96 ± 0.88 (*n* = 13) | 103.98 ± 0.85 (*n* = 16) | 110.60 ± 0.72 (*n* = 19) |
| Stage 1 turning angle (°) | 37.71 ± 29.39 (*n* = 18) | 48.00 ± 49.94 (*n* = 13) | 54.45 ± 44.66 (*n* = 16) | 56.05 ± 33.71 (*n* = 19) |
| Stage 1 turning duration (ms) | 41.20 ± 34.44 (*n* = 18) | 48.08 ± 35.26 (*n* = 13) | 46.88 ± 25.80 (*n* = 16) | 40.79 ± 15.31 (*n* = 19) |
| Stage 1 turning rate (° s<sup>-1</sup>) | 1025.55 ± 354.46 (*n* = 18) | 974.00 ± 469.76 (*n* = 13) | 1171.04 ± 429.53 (*n* = 16) | 1372.06 ± 405.84 (*n* = 19) |
| Stage 2 turning angle (°) | 33.71 ± 9.01 (*n* = 16) | 29.77 ± 9.76 (*n* = 10) | 29.87 ± 10.28 (*n* = 11) | 30.37 ± 11.67 (*n* = 15) |
| Stage 2 turning duration (ms) | 40.36 ± 8.29 (*n* = 16) | 41.25 ± 9.31 (*n* = 10) | 41.67 ± 8.74 (*n* = 11) | 43.33 ± 10.66 (*n* = 15) |
| Stage 2 turning rate (° s<sup>-1</sup>) | 866.18 ± 268.33 (*n* = 16) | 745.13 ± 293.62 (*n* = 10) | 753.74 ± 345.85 (*n* = 11) | 778.11 ± 411.71 (*n* = 15) |
| Apparent looming threshold (rad s<sup>-1</sup>) | 3.55 ± 2.29 (*n* = 18) | 3.90 ± 3.70 (*n* = 12) | 3.63 ± 2.16 (*n* = 15) | 2.68 ± 2.05 (*n* = 19) |
| Maximum acceleration (cm s<sup>-2</sup>) | 6109.23 ± 2156.49 (*n* = 18) | 5659.69 ± 2412.32 (*n* = 13) | 6031.76 ± 2088.32 (*n* = 16) | 6053.00 ± 1736.15 (*n* = 19) |
| Maximum speed (cm s<sup>-1</sup>) | 92.77 ± 28.54 (*n* = 18) | 88.91 ± 33.35 (*n* = 13) | 95.86 ± 30.33 (*n* = 16) | 98.80 ± 29.88 (*n* = 19) |
| Cumulative distance (cm) | 4.19 ± 1.32 (*n* = 18) | 4.08 ± 1.72 (*n* = 13) | 4.40 ± 1.55 (*n* = 16) | 4.66 ± 1.34 (*n* = 19) |

*n*, sample size; “0 ms”, “33 ms”, “83 ms” and “control” show 0 ms treatment, 33 ms treatment, 83 ms treatment, and control, respectively. Escape trajectories are expressed as circular-mean ± standard deviation (s.d.). Others are expressed as linear means ± s.d.

## FIGURE LEGENDS

**Fig. S1.**
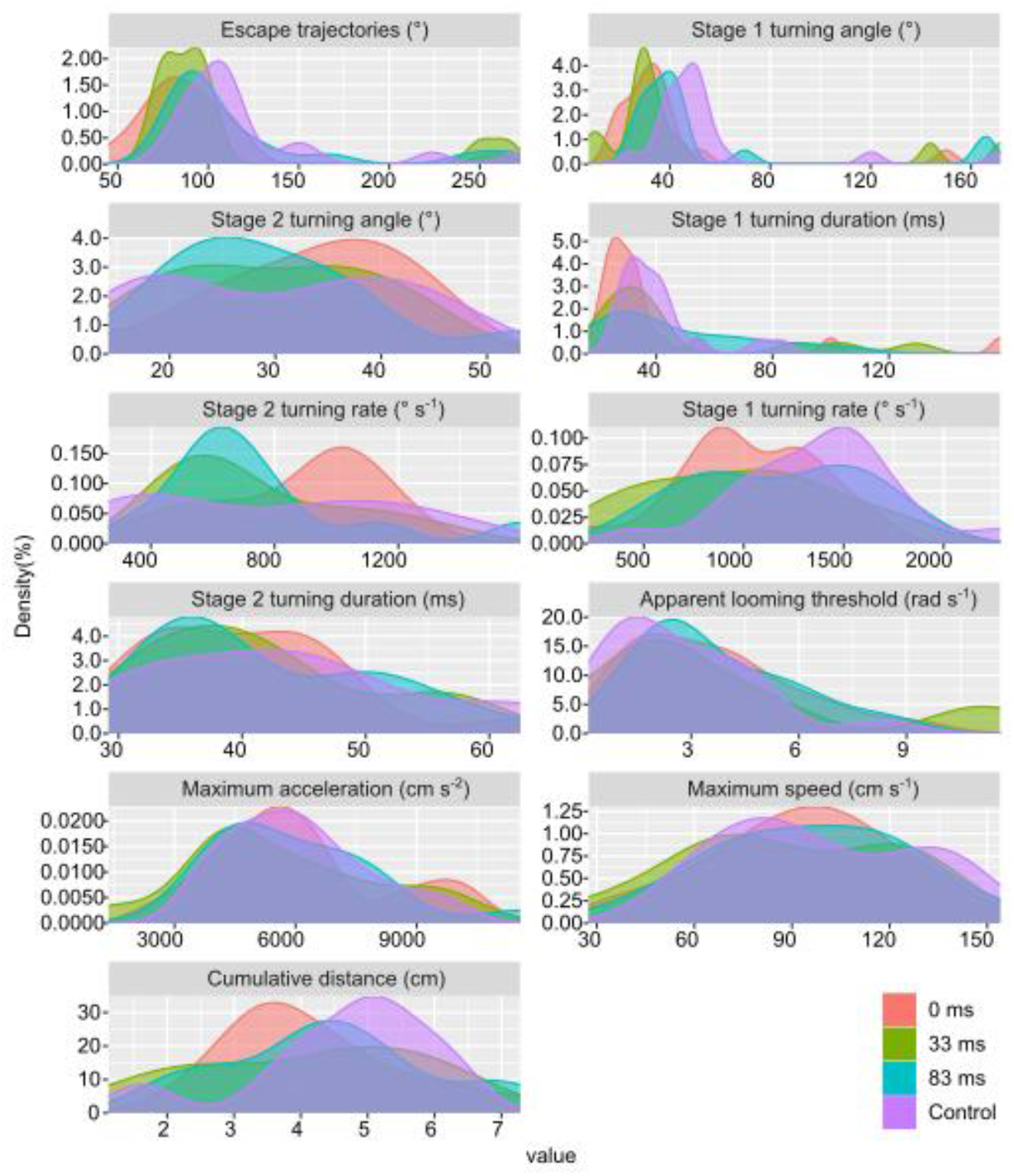
Probability density curve of each variable. Red, green, blue and purple filled curves show 0 ms treatment, 33 ms treatment, 83 ms treatment and control’s data, respectively.

**Fig. S2.**
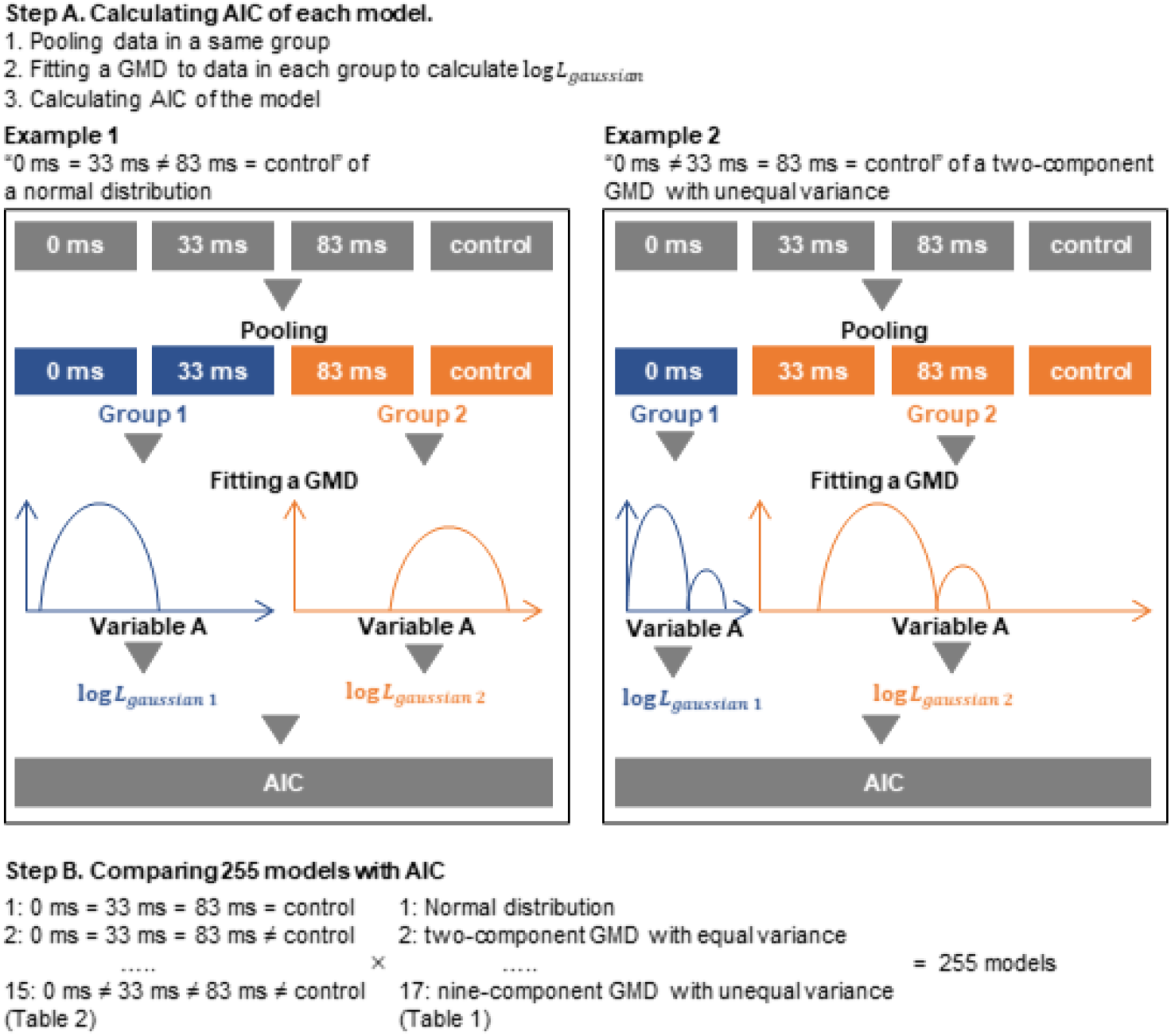
Graphical explanation of the information-theoretic approach to find the best fit model. AIC, Akaike information criterion; GMD, Gaussian mixture distribution; log *L*_*group*_, log-likelihood of each pooled group; “0 ms”, “33 ms”, “83 ms” and “control” show 0 ms treatment, 33 ms treatment, 83 ms treatment, and control, respectively. “=” indicates that there is no different between left and right treatments. “≠” indicates there is a difference between left and right treatments.

